# DIAMOND2GO: A rapid Gene Ontology assignment and enrichment tool for functional genomics

**DOI:** 10.1101/2024.08.19.608700

**Authors:** Christopher Golden, David J. Studholme, Rhys A. Farrer

## Abstract

DIAMOND2GO (D2GO) is a new toolset to rapidly assign Gene Ontology (GO) terms to genes or proteins based on sequence similarity searches. D2GO uses DIAMOND for alignment, which is 100 - 20,000 X faster than BLAST. D2GO leverages GO-terms already assigned to sequences in the NCBI non-redundant database to achieve rapid GO-term assignment on large sets of query sequences. In one test, 98% of the 130,184 predicted human proteins and splice variants were assigned GO-terms (>2 million in total) in < 13 minutes on a laptop computer. D2GO also features the ability to perform enrichment analysis between subsets of data, thereby allowing rapid assignment and detection of over-represented GO-terms in novel sets of sequences. D2GO is freely available under the MIT licence from https://github.com/rhysf/DIAMOND2GO

## Introduction

The Gene Ontology (GO) describes our biological knowledge of molecular function (MF), cellular components (CC) and biological processes (BP). MF describes specific activities performed by gene products, CC describes the locations in the cell where gene products perform their function, and BP describes our understanding of the gene products function [1]. GO-terms use a controlled vocabulary with defined sets of relationships between them and are loosely hierarchical, with ‘child’ terms being more specialized than their ‘parent’ terms. For example, the MF GO:0004375 glycine dehydrogenase (decarboxylating) activity is more specialised than its parent term GO:0003824 catalytic activity. A child term may have several parent terms. GO is developed and maintained by the GO Consortium [1] that runs the GO knowledgebase [2]. The GO Consortium is part of a wider effort by the Open Biological and Biomedical Ontologies (OBO) Foundry [3] to maintain hundreds of separate ontologies across the biological sciences including Cell Ontology, Foundational Model of Anatomy and Plant Ontology [1]. Other examples of functional annotation are Protein Families (Pfam) [4] and Kyoto Encyclopaedia of Genes and Genomes (KEGG) terms [5]. Functional annotation can be assigned to newly annotated genes based on sequence similarity, or other genomic features.

Several tools have been developed to assign GO terms to protein or nucleotide sequences including Blast2GO (B2GO), which is a widely used tool that applies BLAST or the DIAMOND search tool [6] to assign GO terms on the basis of homology to experimentally-characterised proteins [7]. B2GO searches input sequences against either a user defined local database, or a predefined reference database such as the NCBI non-redundant database or UniProtKB/Swiss-Prot [8] via either CloudBlast [9], NCBI QBlast server or Amazon Web Services (AWS) Blast. Assignment of GO-terms is then achieved on the basis of those results along with results of an InterProScan search [10] through an elaborate algorithm that considers similarity, the extension of the query-hit match, the database of choice, the GO hierarchy, and the quality of the original annotations.

B2GO includes a range of additional functionality in addition to GO-term assignment, including visualisation and analysis, and is now part of a larger product named OmicsBox [11] that facilitates various genome and transcriptome analyses with a user-friendly graphical user interface particularly tailored to non-bioinformatics-oriented scientists. By default, it uses one of several online BLAST servers for mapping genes, usually the NCBI non-redundant database of proteins, which can be slow (taking several minutes per batch of queries or individual query sequence). A local copy of the database can be acquired and used, although this requires resources and technical knowledge, and may take considerable time to search depending on the size of the database. With large sets of query sequences, such as the predicted complete proteome from a newly sequenced genome, this initial BLAST step can delay downstream genomic analysis. Furthermore, B2GO requires a subscription or license purchase after the initial (currently) 7-days free trial even for academic users.

Additional promising methods for assigning GO-terms include machine learning approaches trained on additional sources of information such as predicted domains, GO-term frequencies or phylogenetic relationships [12, 13]. GO assignment can also be achieved using EggNog [14] that uses pre-calculated orthology assignments as well as Wei2GO [15] that uses DIAMOND and HMMScan sequence alignment searches against the UniProtKB and Pfam databases. EggNog-Mapper and Wei2GO are both open-access.

The Critical Assessment of Functional Annotation (CAFA) is a community challenge that seeks to evaluate computational annotation of protein function using a time-delayed evaluation method, where new experimentally verified annotations are used as a truth set after results of various tools have been submitted [16]. CAFA3 found a slight improvement of methods between 2016-2019 for MF and BP but not CC. The top-performing CAFA3 method was GOLabeler, which is a machine learning approach trained on several features including GO term frequency, sequence alignment and amino acid trigrams [12]. However, currently the GOLabeler website is not operational, and the software is not publicly available (last access attempt: 19/8/24).

There is a wide range of tools for working with sets of GO terms [17] and genes with assigned GO terms can be used for a variety of investigations. For example, PANTHER can be used to understand the evolutionary and functional classification of protein-coding genes [18]. AmiGO is a web-based tool used for searching, sorting, analysing, and visualizing GO-terms [19]. GO Causal Activity Modelling (GO-CAM) can assist with pathway and network analysis using multiple GO annotations linked into integrated models [20].

In 2014, a new tool named DIAMOND was released that performs pairwise alignment of proteins and translated DNA at 100x-10,000x the speed of BLAST [6]. This could offer a potential time-saving advantage over BLAST-based B2GO. Further improvements to B2GO can be made, such as searching against only the subset of sequences in the database having assigned GO-terms, thus saving time by reducing the search space to less than 1 Gigabases. By combining improvements from the DIAMOND search tool with custom parsing and analysis tools, functional assignment and analysis can be accelerated considerably. Here, we present our new tool, DIAMOND2GO (D2GO), which can assign millions of GO-terms to hundreds of thousands of proteins in minutes on a laptop computer (refer to implementation section).

## Implementation

All analyses in this paper were conducted on a 2021 MacBook Pro with an Apple M1 Max CPU and 64 GB RAM. The NCBI non-redundant database was downloaded on the 14th May 2023 and cleaned for non-printable ASCII characters. NCBI gene2accession and gene2go were downloaded on the 20th July 2023 from https://ftp.ncbi.nih.gov/gene/DATA/. GO-terms and gene accessions were merged using the DIAMOND2GO (D2GO) utility script ncbi_gene2go_merge.pl. GO terms were then added to sequence descriptions using the D2GO utility script blast_database_to_new_description.pl. This new database was indexed using DIAMOND makedb [6]. This database is already included as part of the D2GO Github repository (stored as a Git Large File Storage, which must be installed as a pre-requisite). Instructions to recreate this database are described in **Fig. 1a**.

**Fig. 1.**
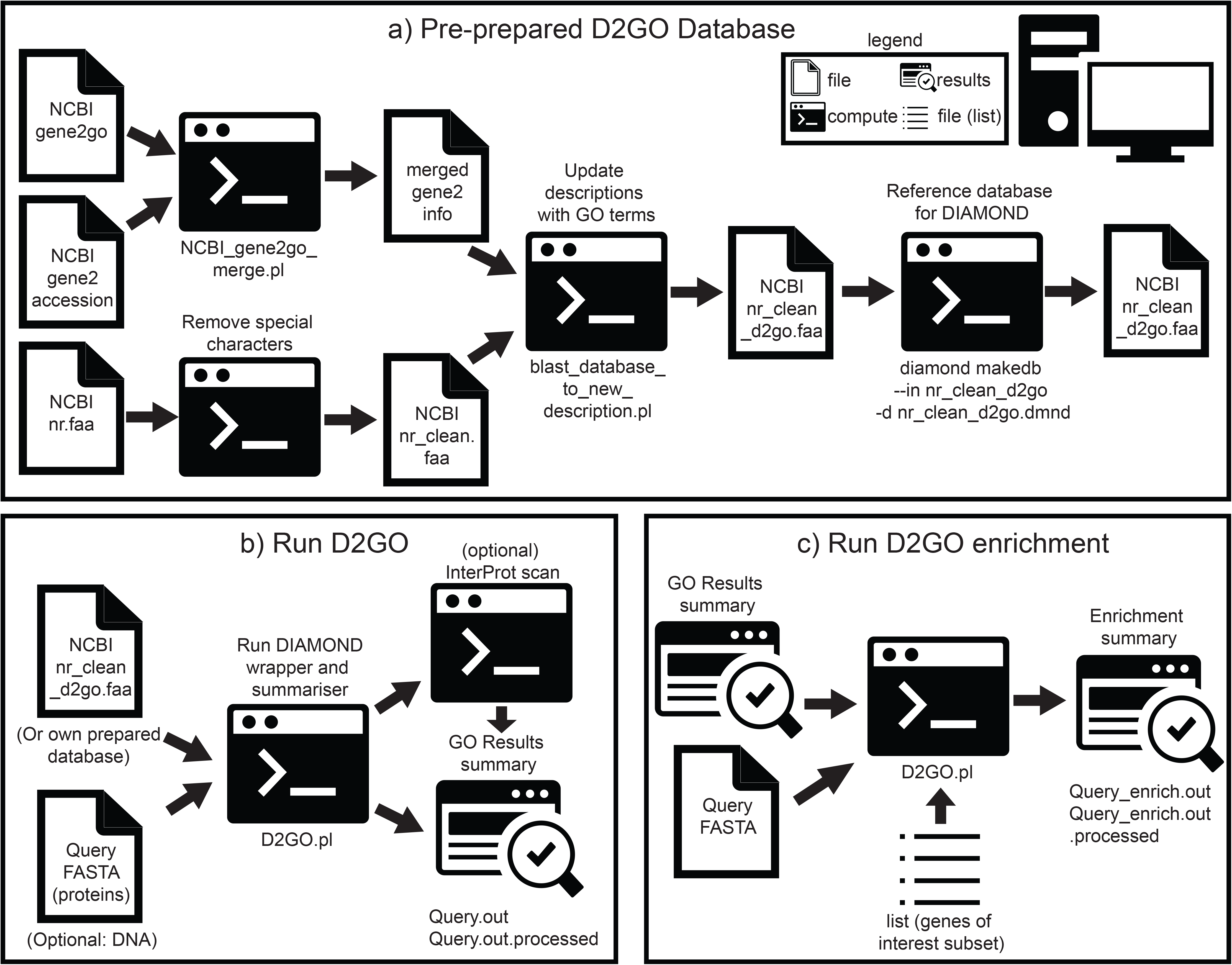
A schematic overview of the steps to construct a new database for Diamond2GO (D2GO), run D2GO and run enrichment analysis. Symbols are described in the embedded legend. All scripts and steps are included as part of the software.

D2GO acts as a wrapper around DIAMOND, including mapping and output parsing functions. Currently either protein (blastp) or nucleotide (blastx) searches are supported, along with sensitivity (default setting: ultra-sensitive), an expectancy value cut-off (default setting: 1e-10), and a max target number of sequences (default setting: 1). A schematic of the pipeline is shown in **Fig. 1b**. The algorithm for this pipeline consists of four steps. First, DIAMOND is run (adjusting parameters described in this paper). The second step summarises the DIAMOND results in a tabulated text file including gene name, species name, and GO-terms and evidence codes. ‘NOT’ Terms (e.g., ‘NOT located_in’) are discarded for brevity. Redundant GO terms for a given gene are removed (keeping only the lowest E-value). The optional third and fourth step is to use the online InterProScan script [10]. The third step prepares the sequences for InterProScan by parsing the query sequences (can be all sequences or only those without DIAMOND hits), removing STOP codons, and splitting the sequences into batches of 500 sequences. The fourth step runs InterProScan, parses the output to tabulated format and combines with the step 1 and step 2 D2GO output.

D2GO was tested on every predicted human protein and splice variant (GRCh38.p14) downloaded from NCBI GenBank (assembly accession GCA_000001405.29) [21, 22] with default sensitivity, 1 max target sequences, and e-value cut-off of 1e^-5^. D2GO was also tested on all predicted proteins of the recently assembled and annotated chytrid fungus *Batrachochytrium salamandrivorans* (*Bsal*) downloaded from NCBI GenBank (assembly accession GCA_002006685.2) [23]. B2GO version 6.0.3 was run on those same *Bsal* proteins as part of that paper, and as a comparison set here [23]. The same *Bsal* proteins were annotated using EggNog v 2.1.12-3 with parameters -m diamond --decorate_gff –excel --cpu 12 -- override. InterProt iprscan5.pl [10] is bundled with D2GO and wrapper scripts are included to run it and compare results. InteractiVenn was used to generate the venn diagram [24]. Significant GO-term enrichment is identified using two-tailed Fisher’s exact test with q-value FDR. Multiple testing corrections were performed with the Storey-Tibshirani [25] method (q-value < 0.05).

## Results

We present DIAMOND2GO (D2GO), which can rapidly assign GO-terms to a query set of sequences (with no subscription or licence costs). We tested D2GO on all 130,184 predicted human proteins, including splice variants, assigning 2,060,956 GO terms to 127,625 proteins (>98% of all proteins). The alignments and output parsing took only 12 minutes and 35 seconds on a laptop computer (refer to implementation section). In contrast, B2GO took several days, while EggNog took 29 minutes and 38 seconds.

To compare D2GO results with other functional annotation assigning tools, we ran D2GO, B2GO and EggNog against the predicted proteins encoded by the chytrid fungus *Bsal* [23] using a 8 different D2GO parameters (**Sup. Table 1**). We then compared the genes with GO terms and the assigned GO terms in each gene across different tools using compare_go_tools.pl (D2GO utility script). D2GO assigned between 68,236 and 203,241 GO-terms (depending on parameters used), which were assigned to between 5,756 and 7,729 genes (53-71% of all 10,867 genes). In comparison, Eggnog assigned 467,171 GO-terms to 4,934 genes (45%), while B2GO assigned 34,408 GO-terms to 8,683 genes (80%). EggNog assigned the highest proportion of GO-terms that were not assigned by other tools (between 88 and 94% of all terms) compared with 45-69% for D2GO and 28-36% for B2GO.

The parameters of D2GO substantially impacted the resulting number of GO-terms assigned and the number of genes with assigned GO-terms. Enforcing a maximum of 25 targets resulted in greater numbers of GO-terms assigned, and a greater proportion that were uniquely assigned compared with other tools, suggesting lower accuracy. Restricting matches to the top hit lowered uniquely assigned GO-terms and therefore comparability with the other tools. As expected, more-stringent thresholds for e-values resulted in fewer genes being assigned GO-terms and a smaller intersection of assigned GO-terms between B2GO and D2GO. Using ultra-sensitive settings resulted in an extra 5-6% of genes being assigned GO-terms. For example, using D2GO parameters e-value < 1E-5 with 1 max target resulted in 57% of genes being assigned GO-terms. Changing the DIAMOND sensitivity from default to ultra increased the number of genes assigned GO-terms to 64%, at the expense of increasing computational time (from < 10 seconds, to ∼9.5 minutes). We have therefore set these settings as default in D2GO as a reasonable trade-off between fast computational time and higher GO-term assignment and show the overlap of genes (g) and terms (t) compared with B2GO and EggNog (**Fig. 2**).

**Fig. 2.**
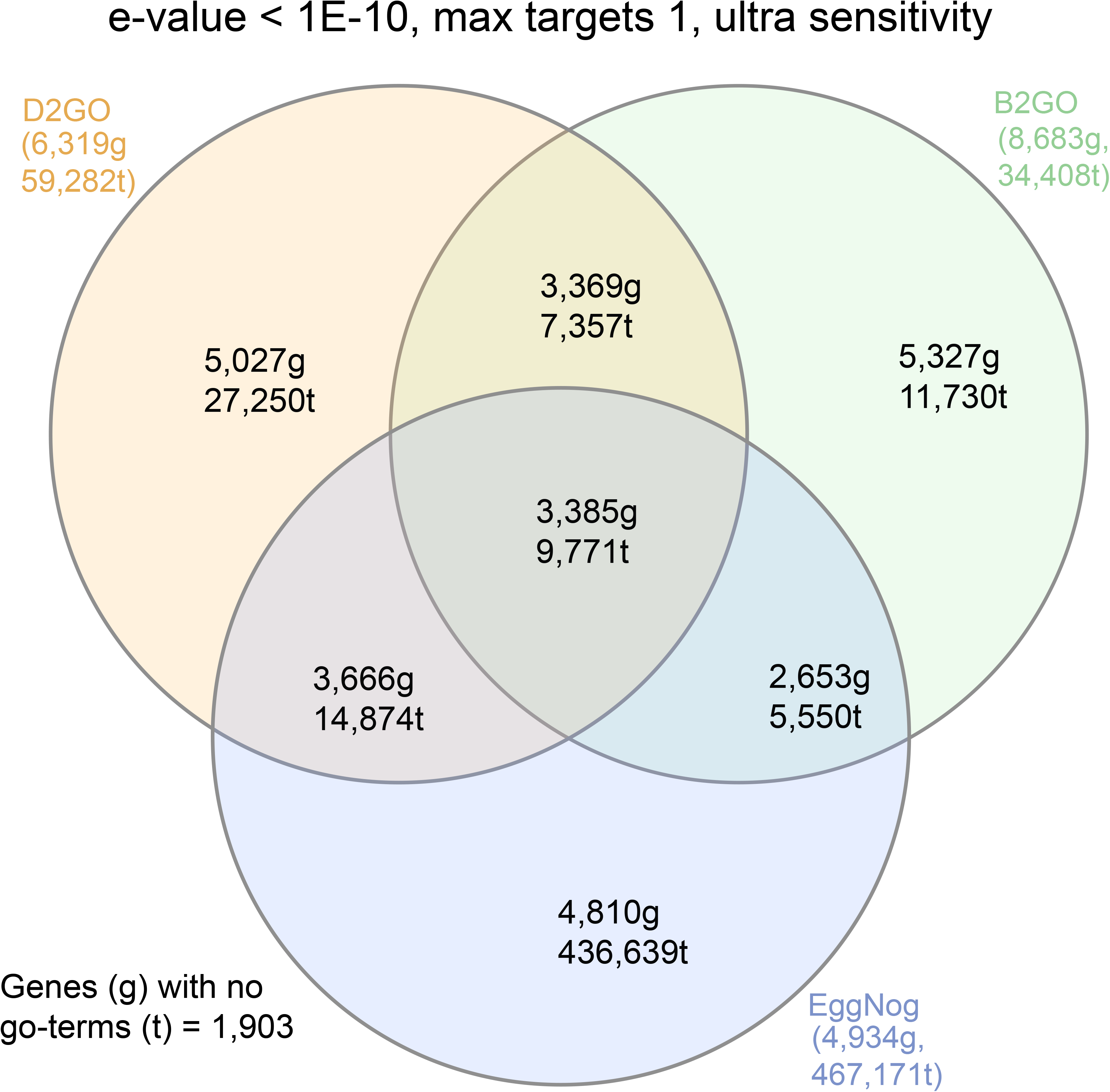
Venn Diagram showing the number of *Bsal* genes (g) that were assigned GO-terms by D2GO (recommended parameters), B2GO and Eggnog. The total number of GO-terms (t) that were assigned to *Bsal* genes by each tool is also shown.

Using D2GO’s inbuilt InterProScan wrapper, a further 7% of genes could be assigned GO-terms (up to 71% of genes). InterPro substantially increased computational time to ∼3 hours when run on genes with no D2GO terms, and ∼17 hours when run on all genes. Running on all genes resulted in only a modest increase in intersect between B2GO and D2GO GO-terms (**Sup. Table 1**). Therefore, using InterProScan loses D2GO competitive GO-term assignment speed, and we do not recommend using those parameters. Together, D2GO has a clear benefit in speed when used without InterProScan (majority of *Bsal* genes assigned GO-terms in < 10 seconds) and shows fair concordance with B2GO and Eggnog assigned GO-terms.

Without a source of ground truth, when faced with discrepancies in GO-term assignment by competing tools, it is impossible to assess their accuracies. An alternative strategy could in future assess D2GO at an upcoming CAFA challenge such as the CAFA5. However, by examining the results for individual well characterised genes, we can quickly assess how reasonable those results look and hypothesise that a similar accuracy is achieved for other less-well-characterised genes. For example, for the DNA-directed DNA polymerase alpha subunit pol12 (locus ID BSLG_001742), both tools identified the same 3 relevant GO terms, while D2GO (with InterProScan setting turned on) identified 2 additional GO-terms (1 BP and 1 CC) that are parents of 2 of the other terms (**Sup. Table 2**). B2GO BLASTp matched a separate gene in *Bsal* in its database (BSLG_01791) to assign GO terms. In contrast, *Bsal* is absent from the D2GO database (due to no *Bsal* entries currently in the NCBI gene2go file), and instead BSLG_01791 top hit was to *Xenopus laevis* (African clawed frog). Therefore, despite very different gene matches in the database, a complete overlap of GO-terms was recovered by both tools for this gene. These results indicate that D2GO yields similar results to B2GO for at least some genes and does so much faster.

The D2GO utility script ‘test_enrichment.pl’ was run on all human genes with ‘polymerase’ in their FASTA description (**Fig. 1c**). The top 10 most significantly enriched GO-terms were polymerase-related terms as expected (**Sup. Table 3**), suggesting that the GO-terms assigned by D2GO and the enrichment tests will be useful for less-well characterised gene categories.

In conclusion, D2GO offers an alternative to other functional annotation tools, such as B2GO and Eggnog. D2GO is quicker than both tools, and freely available like Eggnog.

## Legends

**Sup. Table 1**. GO-terms assigned to *Bsal* proteins by D2GO using different settings, with or without InterPro (N=no, no_hits=run on genes with no DIAMOND hits, all=run on all genes). Overlap with B2GO is presented for comparison.

**Sup. Table 2**. GO-terms identified for the DNA-directed DNA polymerase alpha subunit pol12 (locus ID BSLG_001742) in *Bsal* by B2GO and D2GO (using InterProScan on all genes). Go-term sub-ontologies are Biological Process (BP), Cellular Component (CC) and Molecular Function (MF). Parental terms of other GO-terms identified for this gene are shown.

**Sup. Table 3**. Top 10 significant GO-terms for Human genes that include the word ‘polymerase’ in the description.

## Acknowledgements

We’d like to thank Chris Desjardins for Perl code involved in testing enrichment.

## Declarations

### Ethics approval and consent to participate

Not applicable

### Consent for publication

Not applicable

### Availability of data and materials

**Project name:** DIAMOND2GO

**Project home page**: https://github.com/rhysf/DIAMOND2GO

**Operating system(s):** Platform independent. Tested on MacOS.

**Programming language:** Perl

**Other requirements:** Git Large File Storage, BioPerl, CPAN modules Getopt::Std and Scalar::Util, DIAMOND.

**License:** MIT license.

**Any restrictions to use by non-academics:** No

### Authors’ contributions

CG, DJS and RAF wrote the main manuscript and prepared figures and tables. CG and RAF wrote the software. All authors reviewed the manuscript.

## References

1. Ashburner M, Ball CA, Blake JA, Botstein D, Butler H, Cherry JM, et al. Gene Ontology: tool for the unification of biology. Nat Genet. 2000;25:25–9.

2. The Gene Ontology Consortium, Aleksander SA, Balhoff J, Carbon S, Cherry JM, Drabkin HJ, et al. The Gene Ontology knowledgebase in 2023. Genetics. 2023;224:iyad031.

3. Smith B, Ashburner M, Rosse C, Bard J, Bug W, Ceusters W, et al. The OBO Foundry: coordinated evolution of ontologies to support biomedical data integration. Nat Biotechnol. 2007;25:1251.

4. Finn RD, Bateman A, Clements J, Coggill P, Eberhardt RY, Eddy SR, et al. Pfam: the protein families database. Nucleic Acids Res. 2014;42 Database issue:D222–30.

5. Kanehisa M, Goto S. KEGG: kyoto encyclopedia of genes and genomes. Nucleic Acids Res. 2000;28:27–30.

6. Buchfink B, Xie C, Huson DH. Fast and sensitive protein alignment using DIAMOND. Nat Methods. 2015;12:59–60.

7. Conesa A, Götz S, García-Gómez JM, Terol J, Talón M, Robles M. Blast2GO: a universal tool for annotation, visualization and analysis in functional genomics research. Bioinformatics. 2005;21:3674–6.

8. Bairoch A, Apweiler R. The SWISS-PROT protein sequence database and its supplement TrEMBL in 2000. Nucleic Acids Res. 2000;28:45–8.

9. Matsunaga A, Tsugawa M, Fortes J. CloudBLAST: Combining MapReduce and Virtualization on Distributed Resources for Bioinformatics Applications. In: 2008 IEEE Fourth International Conference on eScience. 2008. p. 222–9.

10. Hunter S, Apweiler R, Attwood TK, Bairoch A, Bateman A, Binns D, et al. InterPro: the integrative protein signature database. Nucleic Acids Res. 2009;37 Database issue:D211–5.

11. Bioinformatics Software OmicsBox. BioBam. https://www.biobam.com/omicsbox/. xAccessed 1 Feb 2024.

12. You R, Zhang Z, Xiong Y, Sun F, Mamitsuka H, Zhu S. GOLabeler: improving sequence-based large-scale protein function prediction by learning to rank. Bioinformatics (Oxford, England). 2018;34.

13. Fa R, Cozzetto D, Wan C, Jones DT. Predicting human protein function with multi-task deep neural networks. PLOS ONE. 2018;13:e0198216.

14. Huerta-Cepas J, Forslund K, Coelho LP, Szklarczyk D, Jensen LJ, von Mering C, et al. Fast Genome-Wide Functional Annotation through Orthology Assignment by eggNOG-Mapper. Molecular Biology and Evolution. 2017;34:2115–22.

15. Reijnders MJMF. Wei2GO: weighted sequence similarity-based protein function prediction. PeerJ. 2022;10:e12931.

16. Zhou N, Jiang Y, Bergquist TR, Lee AJ, Kacsoh BZ, Crocker AW, et al. The CAFA challenge reports improved protein function prediction and new functional annotations for hundreds of genes through experimental screens. Genome Biology. 2019;20:244.

17. Shahzad M, Ahsan K, Nadeem A, Sarim M. Gene Ontology Tools: A Comparative Study. J basic appl sci. 2015;11:619–29.

18. Thomas PD, Ebert D, Muruganujan A, Mushayahama T, Albou L-P, Mi H. PANTHER: Making genome-scale phylogenetics accessible to all. Protein Science. 2022;31:8–22.

19. Carbon S, Ireland A, Mungall CJ, Shu S, Marshall B, Lewis S, et al. AmiGO: online access to ontology and annotation data. Bioinformatics. 2009;25:288–9.

20. Thomas PD, Hill DP, Mi H, Osumi-Sutherland D, Van Auken K, Carbon S, et al. Gene Ontology Causal Activity Modeling (GO-CAM) moves beyond GO annotations to structured descriptions of biological functions and systems. Nat Genet. 2019;51:1429–33.

21. Lander ES, Linton LM, Birren B, Nusbaum C, Zody MC, Baldwin J, et al. Initial sequencing and analysis of the human genome. Nature. 2001;409:860–921.

22. International Human Genome Sequencing Consortium. Finishing the euchromatic sequence of the human genome. Nature. 2004;431:931–45.

23. Wacker T, Helmstetter N, Wilson D, Fisher MC, Studholme DJ, Farrer RA. Two-speed genome evolution drives pathogenicity in fungal pathogens of animals. Proc Natl Acad Sci U S A. 120:e2212633120.

24. Heberle H, Meirelles GV, da Silva FR, Telles GP, Minghim R. InteractiVenn: a web-based tool for the analysis of sets through Venn diagrams. BMC Bioinformatics. 2015;16:169.

25. Storey JD, Tibshirani R. Statistical significance for genomewide studies. Proc Natl Acad Sci USA. 2003;100:9440–5.

